## Supplementary Material for "Folding correctors can restore CFTR posttranslational folding landscape by allosteric domain-domain coupling"

### RESOURCE AVAILABILITY

#### Lead contact

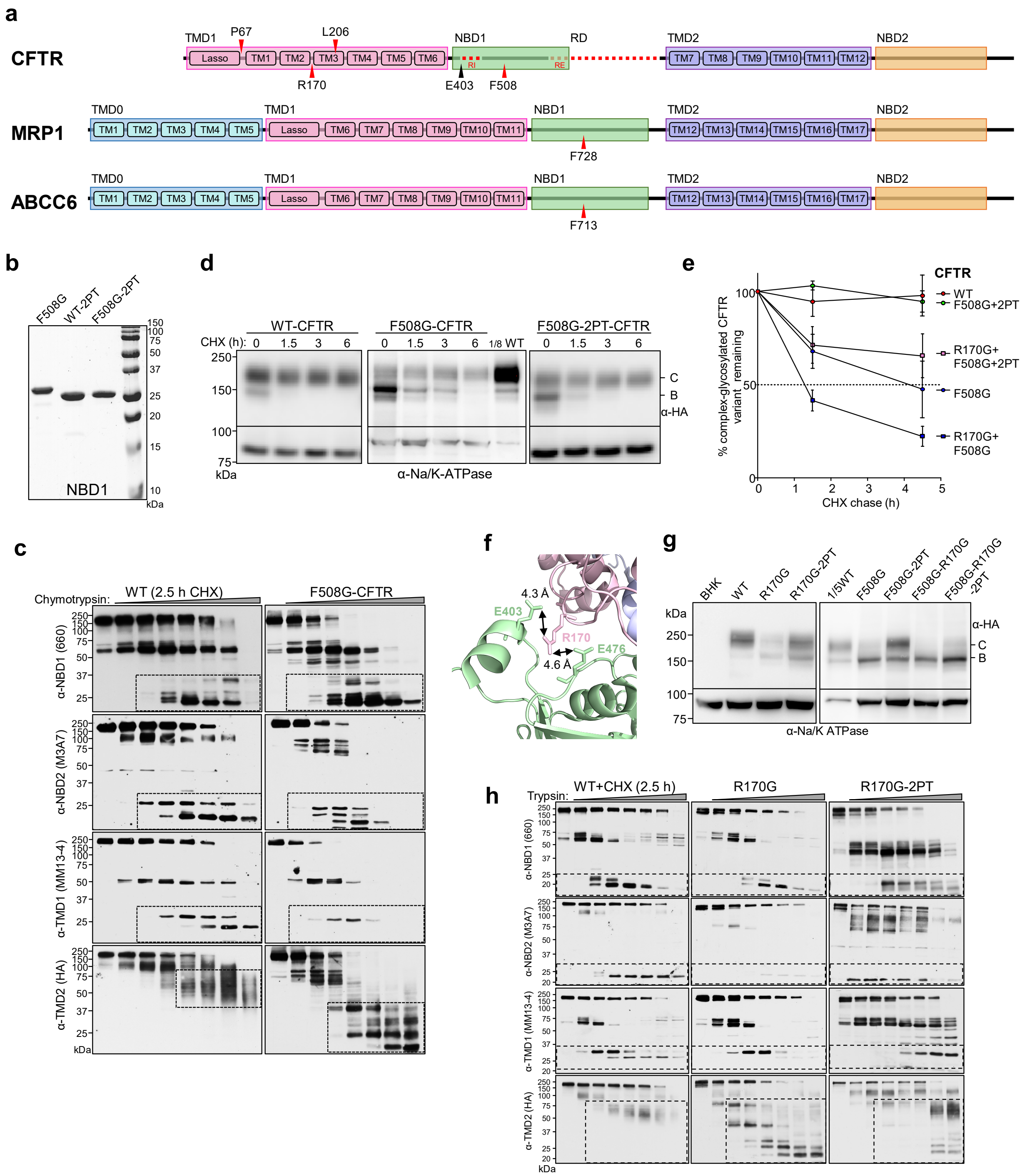


**Supplementary Figure 1. Uncoupling of NBD1-CL1(TMD1) or NBD1-CL4(TMD2) interfaces causes defects in CFTR in vivo biogenesis, conformational and metabolic stability, which can be reversed suppressor mutation.**

**a)** Mutations introduced into the indicated domains of ABCC transporters. The color-coding of CFTR, MRP1, and ABCC6 domains follows as defined in Fig. 1a left panel. The TMD0s of MRP1 and

ABCC6 are in cyan. **b)** Purified CFTR NBD1 variants in the absence and presence of 2PT suppressor mutations were visualized by SDS-PAGE and Coomassie Blue staining. **c)** The conformational stabilities of WT- and F508G-CFTR were determined with limited chymotrypsin digestion (at 0 °C) in isolated microsomes and then probed with immunoblotting, using domain specific antibodies. **d-e)** The metabolic stability of the complex-glycosylated WT-, F508G- and F508G-2PT-CFTR was measured after translational inhibition with cycloheximide (CHX) for the indicated time. CFTR abundance was measured in equal amounts of cell lysates (except for the WT-CFTR) by quantitative immunoblotting. Means ± S.E.M., n = 4, biological replicates. **f)** Electrostatic interaction of the R170 residue (CL1) with the E403 and E476 of NBD1 in the human CFTR cryo-EM structure (PDB:6MSM). **g)** Comparison of the steady-state expression levels of complex-glycosylated (C-band) CFTR variants in stably transfected BHK-21 cell lysates by immunoblotting. Equal amounts of lysates (50 μg) were probed, but for WT-CFTR (10 μg), as indicated. Representative of three experiments. The 2PT-suppressor rescue efficiency on the complex-glycosylated R170G- and F508G-CFTR variants was determined quantitative immunoblotting (see Fig.1c). **h)** The TMD1, NBD1, TMD2 and NBD2 conformational stability of R170G- and R170G-2PT-CFTR was probed by limited trypsinolysis and immunoblotting of isolated microsomes by domain specific antibodies. To eliminate the core-glycosylated WT-CFTR, microsomes were isolated from BHK-21 cells after 2.5h treatment with 150 μg/ml CHX. Trypsin concentration used: 0, 1, 3, 10, 30 100, 300, and 1000 μg/ml. Representative of 2-3 experiments. Source data are provided as a Source Data file.

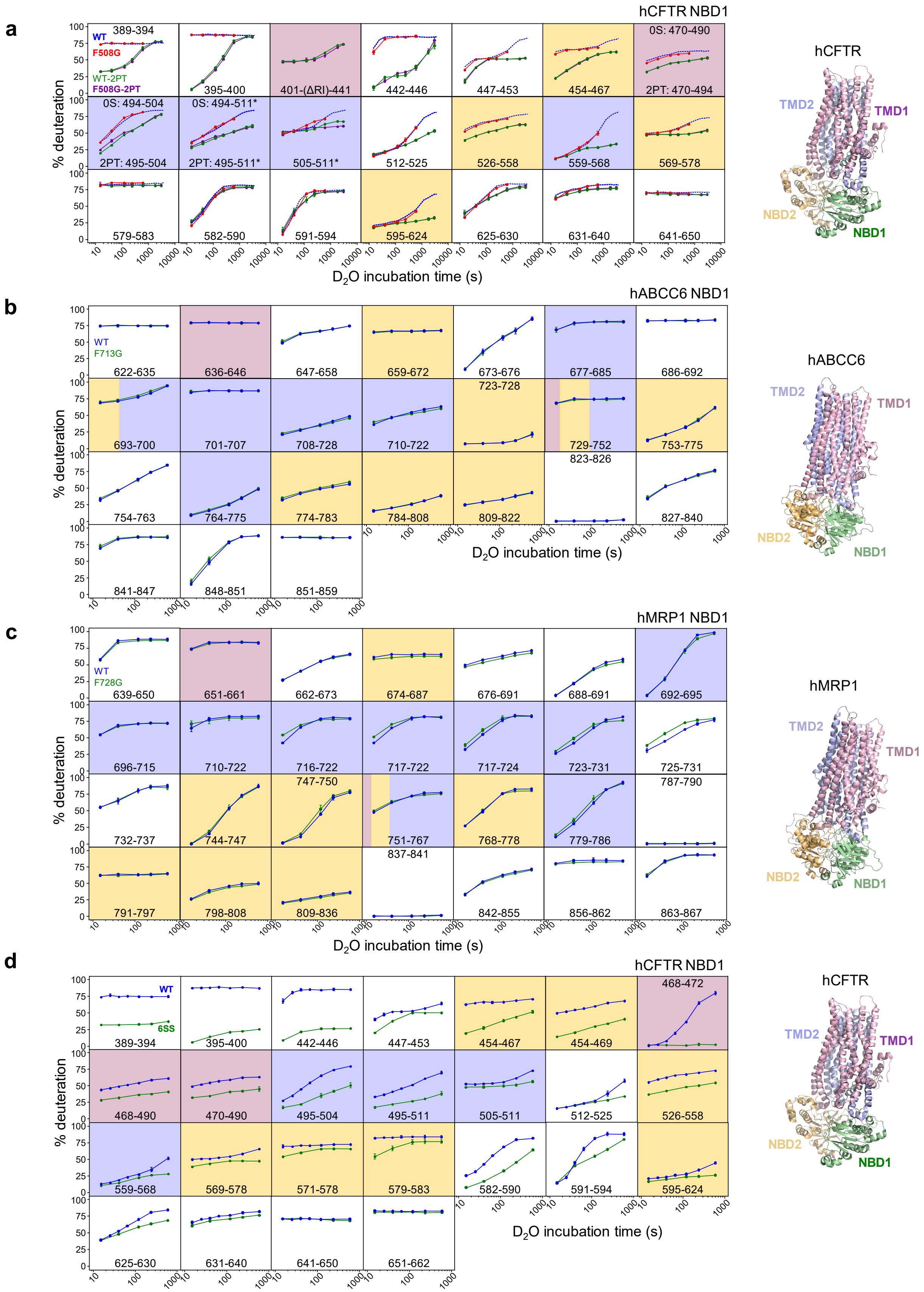
a
hCFTR NBD1
WT
F508G
WT-2PT
F508G-2PT
389-394
395-400
401-(ΔRI)-441
442-446
447-453
454-467
0S: 470-490
2PT: 470-494
0S: 494-504
2PT: 495-504
0S: 494-511*
2PT: 495-511*
505-511*
512-525
526-558
559-568
569-578
579-583
582-590
591-594
595-624
625-630
631-640
641-650
% deuteration
$D_2O$ incubation time (s)
hCFTR
TMD2
TMD1
NBD2
NBD1
b
hABCC6 NBD1
WT
F713G
622-635
636-646
647-658
659-672
673-676
677-685
686-692
693-700
701-707
708-728
710-722
723-728
729-752
753-775
754-763
764-775
774-783
784-808
809-822
823-826
827-840
841-847
848-851
851-859
% deuteration
$D_2O$ incubation time (s)
hABCC6
TMD2
TMD1
NBD2
NBD1
c
hMRP1 NBD1
WT
F728G
639-650
651-661
662-673
674-687
676-691
688-691
692-695
696-715
710-722
716-722
717-722
717-724
723-731
725-731
732-737
744-747
747-750
751-767
768-778
779-786
787-790
791-797
798-808
809-836
837-841
842-855
856-862
863-867
% deuteration
$D_2O$ incubation time (s)
hMRP1
TMD2
TMD1
NBD2
NBD1
d
hCFTR NBD1
WT
6SS
389-394
395-400
442-446
447-453
454-467
454-469
468-472
468-490
470-490
495-504
495-511
505-511
512-525
526-558
559-568
569-578
571-578
579-583
582-590
591-594
595-624
625-630
631-640
641-650
651-662
% deuteration
$D_2O$ incubation time (s)
hCFTR
TMD2
TMD1
NBD2
NBD1

**Supplementary Figure 2. Comparison of isolated WT and mutant NBD1s conformational dynamics of CFTR, MRP1, and ABCC6.**

The time course of deuteration of **a)** WT- and F508G-NBD1 in the absence or presence of 2PT-suppressor mutation. NBD1 peptides that interface the TMD1, TMD2, or NBD2 have pink, light blue and light orange background color, respectively, as the respective domains of the hCFTR cryo-EM structure in Fig.1a. Peptide interactions with more than one domain interface are indicated by multiple colors. The NBD1 inter-domain interfaces were determined using the outward-facing conformation of hCFTR (PDB: 6MSM). **b-c)** The deuteration time course of ABCC6 WT- and F713G-NBD1, as well as MRP1 WT- and F728G-NBD1 was measured by continuous HDX-MS at 37 °C. The NBD1 inter-domain interfaces were determined using the outward-facing conformations of ABCC6 and MRP1 model structures. The interfaces are colored like hCFTR domain-assignment as above and indicated on the homology models of ABCC6 and MRP1 at the right. **d)** WT- and 6SS-NBD1 deuteration kinetics was determined by HDX-MS at 37°C, performing pepsin digestion at 0°C. Means ± S.D., n = 3, technical replicates. The deuteration kinetics of CFTR NBD1 interfaces are shown in the main figures as well (Fig. 2j, 4a). Source data are provided as a Source Data file.

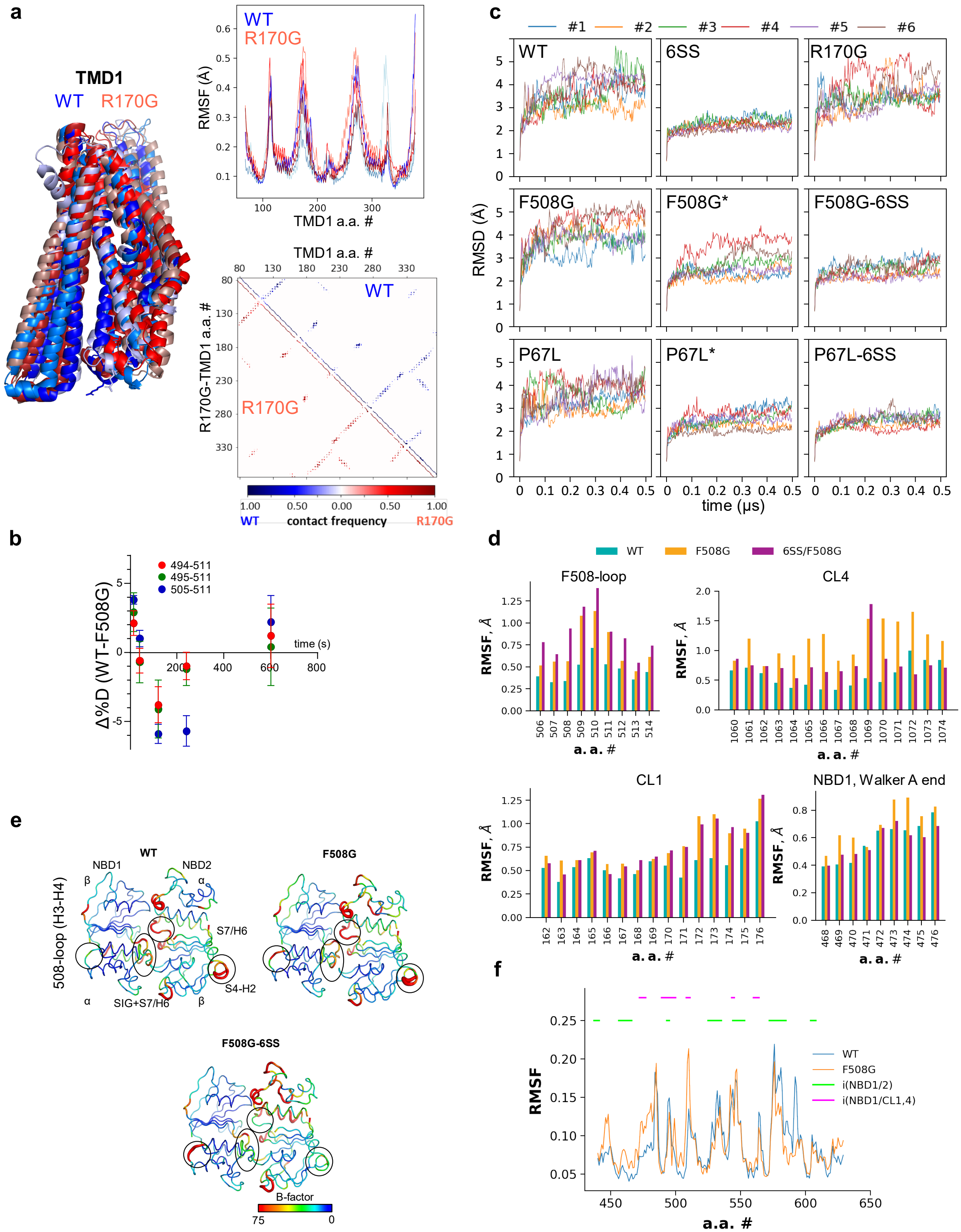


**Supplementary Figure 3. MD simulations and backbone NHs deuteriation kinetics of NBD1-CL4(TMD2) interface uncoupling effect on CFTR domain interfaces and the impact of NBD1 suppressors.**

**a)** MD simulations on isolated WT- (blue) and R170G-TMD1 (red) to assess the effect of R170G mutation on TMD1 dynamics (500 ns, n = 3-3 for each construct, frames = 15,000). No differences in the final conformations were observed (left panel). Quantitative measures, such as RMSF (root mean square fluctuations) of residues (right top panel) and residue-residue contacts (right lower panel) did not reveal considerable structural differences between WT- and R170G-TMD1. **b)** Differential deuteriation of F508-loop peptides in WT- and F508G-NBD1 ($\Delta\%D_{(WT\text{-}F508G)}$) as a function of $D_2O$ incubation. Means ± S.D., n = 3, technical replicates. **c)** The Cα RMSD of each frame from the initial structure was calculated and indicated a sufficient equilibration of MD simulations with the full-length CFTR variants. RMSD values were also calculated from simulations with complete CFTR structures but computed without taking the regulatory insertion (RI) into account ("*" marks) to demonstrate that the high RMSD were caused by the dynamic RI loop, providing better comparison with 6SS constructs lacking the RI. We selected the outward-facing conformation (PDB:6MSM) for simulations with bound ATP molecules, since the NBDs of the inward-facing conformation has been shown to exhibit a high level of rigid body motions[1,2]. The distances between NBD1/2 throughout our simulations are presented in Supplementary Table 3. Furthermore, the non-phosphorylated CFTR structure may be distinct from the physiological conformational ensembles, because it was determined in the absence of ATP. The degenerate ATP-binding Site-1 is occupied by an ATP[3] at cytosolic ATP concentration, thus NBD1/NBD2 remain, predominantly, associated at Site 1. **d)** RMSF was calculated for the F508-loop, CL4, CL1, and Walker A helix for WT-, F508G-, and F508G-6SS-CFTR from six-six simulations (frames = 30,000 each). The RMSF of G1069 in F508G-6SS-CFTR is high and most likely allowed by unrestricted rotations around Cα because of the absence of the Gly side chain. **e)** Dynamics of the NBD1/NBD2 is presented in structural context. Warmer colors and thicker representation depict higher dynamic fluctuations. Largest changes are highlighted by circles. **f)** NBD1 RMSF from WT- and F508G-CFTR simulations were compared. The NBD1 domain interfaces are indicated. Source data are provided as a Source Data file.

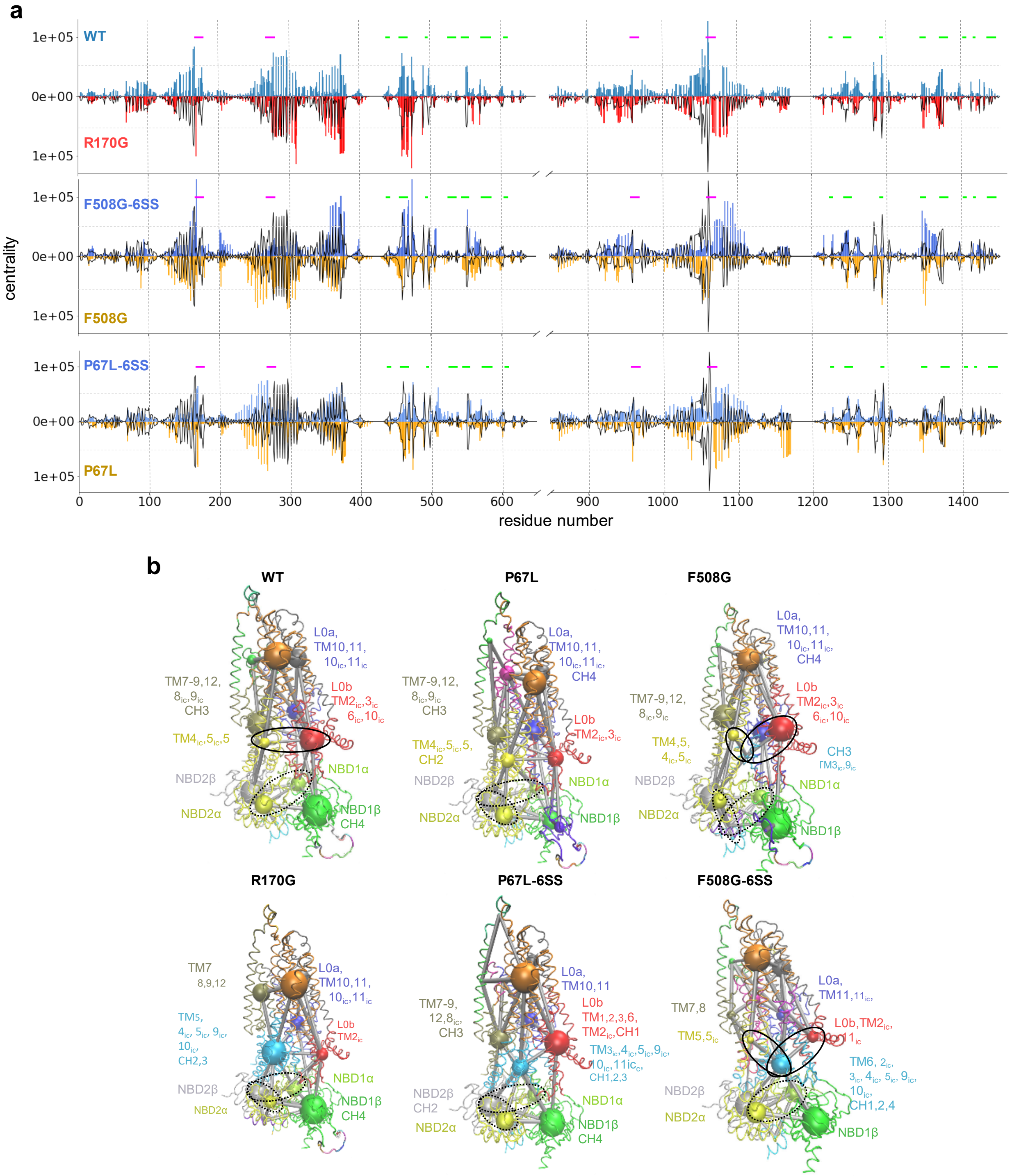


**Supplementary Figure 4. The effect of 6SS-suppressors on allosteric networks and dynamic communities of CFTR variants.**

**a)** To reveal perturbations in propagation of dynamic changes caused by mutations, we built a residue-network, based on pairwise residue motion-correlation as observed in MD simulations for the WT and mutant CFTRs. Betweenness centrality of each node/residue was calculated and plotted to indicate the number of the shortest (allosteric) pathways going through each residue. Distinct rewiring of the allosteric networks was instigated by the R170G, F508G and P67L mutations, based on the differences observed in their centrality patterns. Accordingly, the 6SS-induced conformational rescue was associated with different centrality patterns from that of the WT. For comparison, black lines indicate the WT-CFTR values on each graph. Horizontal lines indicate the position of coupling helices (magenta) and NBD1/NBD2 interface residues (green). **b)** Dynamic communities were depicted in structural context in the indicated CFTR variants. Communities (amino acids in a community) were color-coded. Smaller letters indicate that only a smaller fraction of the region contributes to the community. The spheres indicate the community size. The thickness (1-weight) of edges between spheres represents the dynamic coupling strength between communities. Black dotted circles in the R170G-CFTR highlight allosteric decoupling of NBD1 and NBD2 α-subdomains. Similarly, black dotted circles enclose a path from NBD1 α-subdomain community to NBD2 α-subdomain community to overcome the direct step between them disrupted by the P67L mutation. This path was shortened by 6SS, indicated by the smaller distance between these communities in the P67L-6SS-CFTR, when compared to the P67L-CFTR (0.0408 versus 0.0507; -log($C_{ij}$) values). F508G and F508G-6SS communities are shown for comparison. Black circles highlight the alterations in coupling between the distant TM4/5 and L0/Lasso. Detailed lists of residues in each community and visualization scripts for VMD can be downloaded from https://doi.org/10.5281/zenodo.8388593. Source data are provided as a Source Data file.

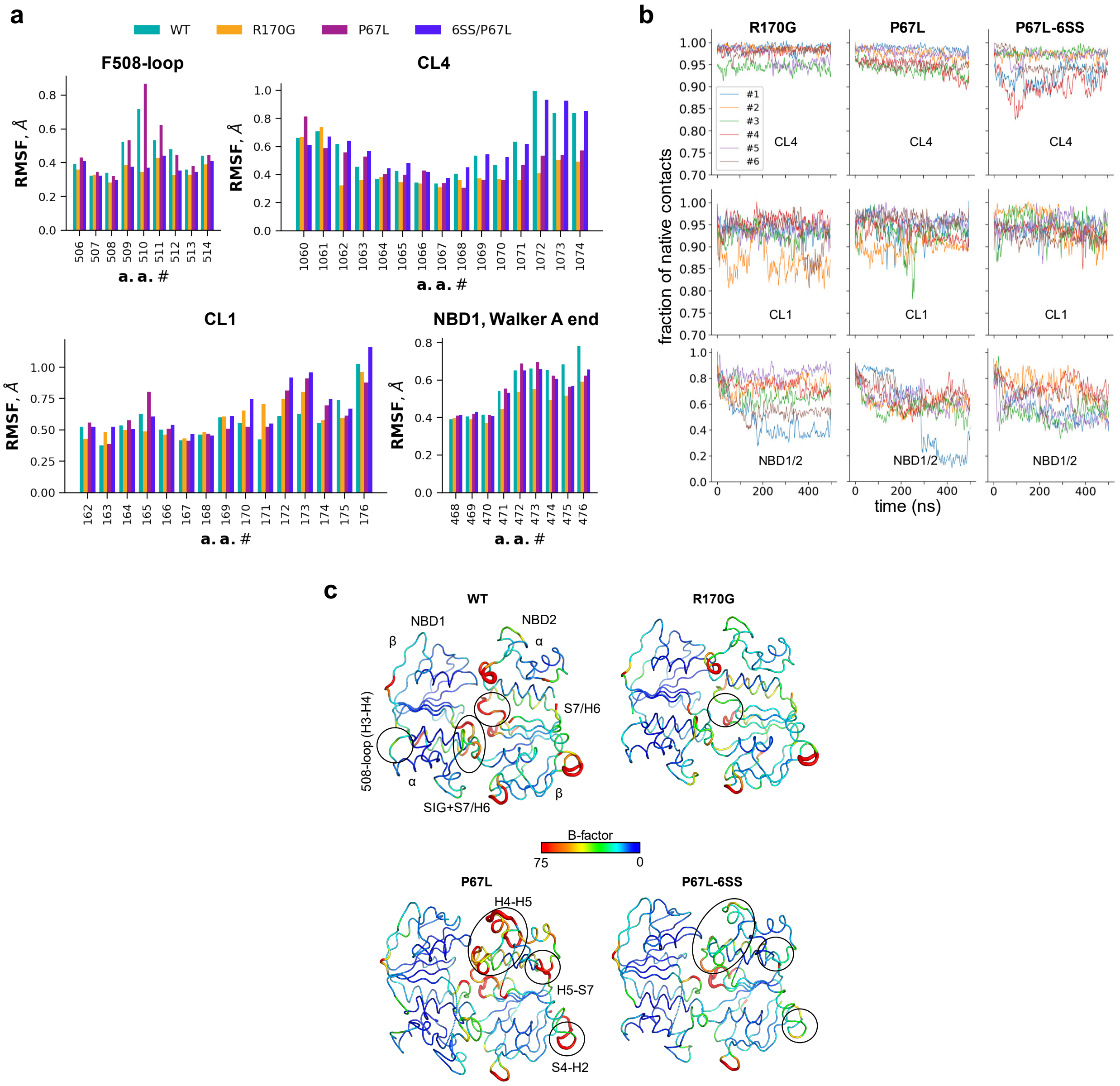


**Supplementary Figure 5. The effect of R170G, P67L, and 6SS suppressors on CFTR's domain contacts and dynamics.**

**a)** RMSF was calculated for the F508-loop, CL4, CL1 and Walker A helix from 6-6 simulations (n = 6-6, frames = 30,000) and is plotted for WT-, R170G-, P67L- and P67L-6SS-CFTR. **b)** Fraction of native contacts of CL4 and CL1 with the rest of the protein and between NBD1 and NBD2 were calculated over all trajectories and plotted for R170G-, P67L- and P67L-6SS-CFTR. **c)** Dynamics of the NBD1/NBD2 is presented in structural context based on MD simulations with WT-, R170G-, P67L- and P67L-6SS-CFTR, calculated using the gmx rmsf tool (frames = 30,000). Warmer colors and thicker representation indicate higher dynamic fluctuations. Differences are highlighted by circles. Source data are provided as a Source Data file.

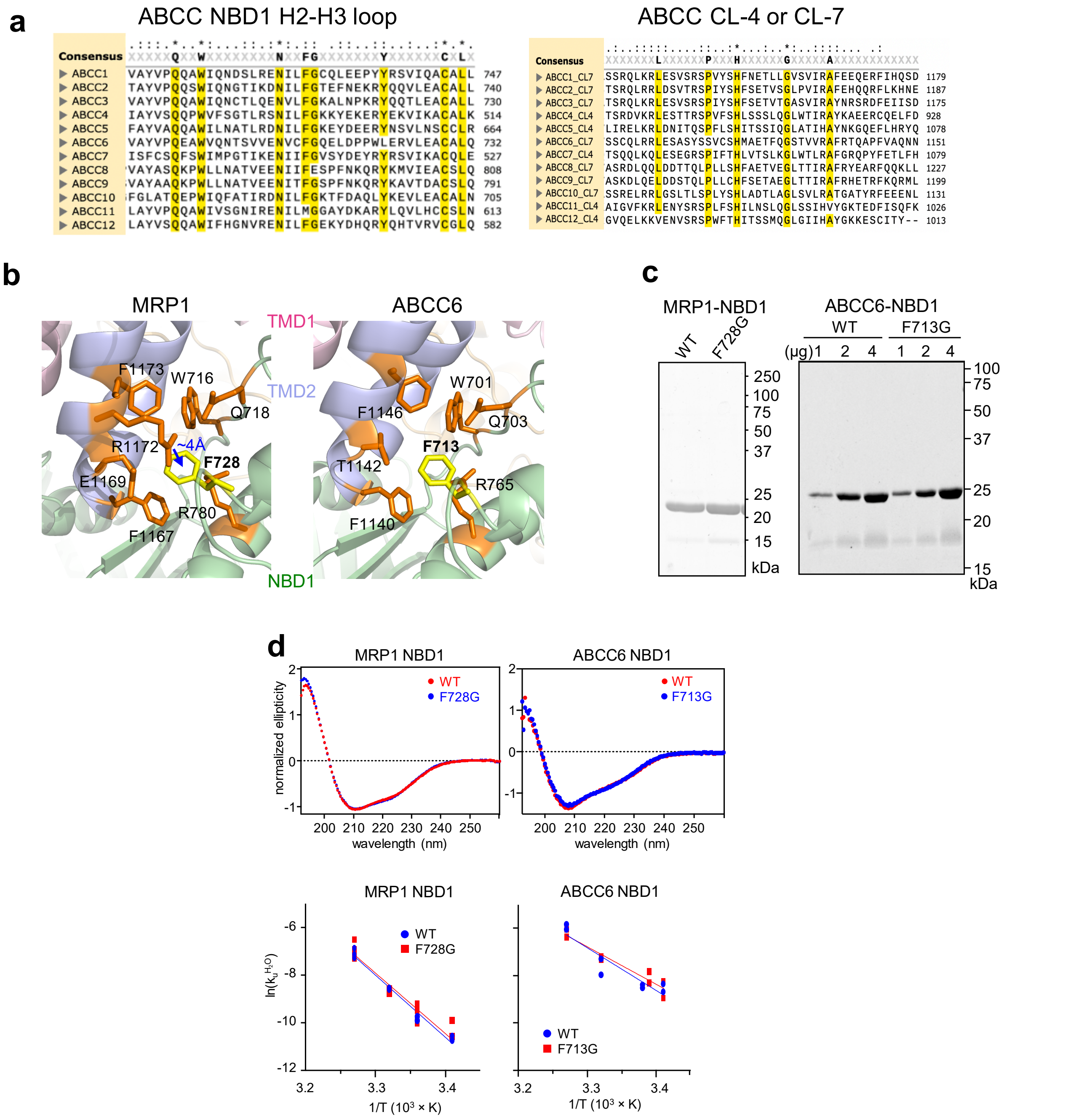


**Supplementary Figure 6. Purification and biophysical characterization of isolated NBD1 variants of human MRP1 and ABCC6.**

**a)** Multiple sequence alignments of human ABCC-transporters' NBD1-TMD2 interface residues, consisting of the NBD1 Phe-loop of H2-H3 (left panel) and the corresponding interface residues of the CL4 or CL7 loop, were generated using Clustal Omega[4]. Conserved residues are labelled as follow: * : fully conserved residues have a conservation scoring (CS) of 1 (yellow), : : CS > 0.5, and . : CS =

0.5 - 0. CS for the R1173 (highlighted in red) in the hMRP1 is 0 - 0.5. **b)** The environment of the F728 and F713 NBD1 residues in the 3D-structures of hMRP1 and hABCC6, respectively, based on homology modelling as described in Methods. The F728 side-chain forms a cation-π interaction with R1173 (CL7/TMD2) in the MRP1, while the F713 may interface a hydrophobic cavity partly consisting of the CL7(TMD2) in ABCC6. Color-coding for NBDs and TMDs of MRP1 and ABCC6 follows that of the hCFTR (Fig.1a). **c)** SDS-PAGE of purified NBD1s of MRP1 and ABCC6 variants. **d)** Top panels: CD spectra for the indicated MRP1 and ABCC6 NBD1 variants. Data are normalized for the ellipticity value at 214 nm and representative of 2-3 measurements. Lower panels: Unfolding rates ($k_u^{H_20}$) of MRP1 and ABCC6 WT- and mutant-NBD1s were determined from the initial loss of ellipticity at 0 - 6M urea concentration (20, 24, 28 and 32 °C) by CD spectroscopy. The $k_u^{H_20}$ values were calculated by linear extrapolation for 37°C. The predicted unfolding activation energy ($\Delta G^{\#}{}_u$) of the WT- and F728G-MRP1 NBD1 at 37 °C were estimated using the extrapolated $k_u^{H_20}$ and were: 3.36 kcal/mol and 3.57 kcal/mol, respectively. The $\Delta G_u^{\#}$ of the WT- and F713G-ABCC6 NBD1 at 37 °C were: 3.26 kcal/mol and 3.35 kcal/mol, respectively. Data are derived from two biological and two technical replicate measurements at each temperature. Source data are provided as a Source Data file.

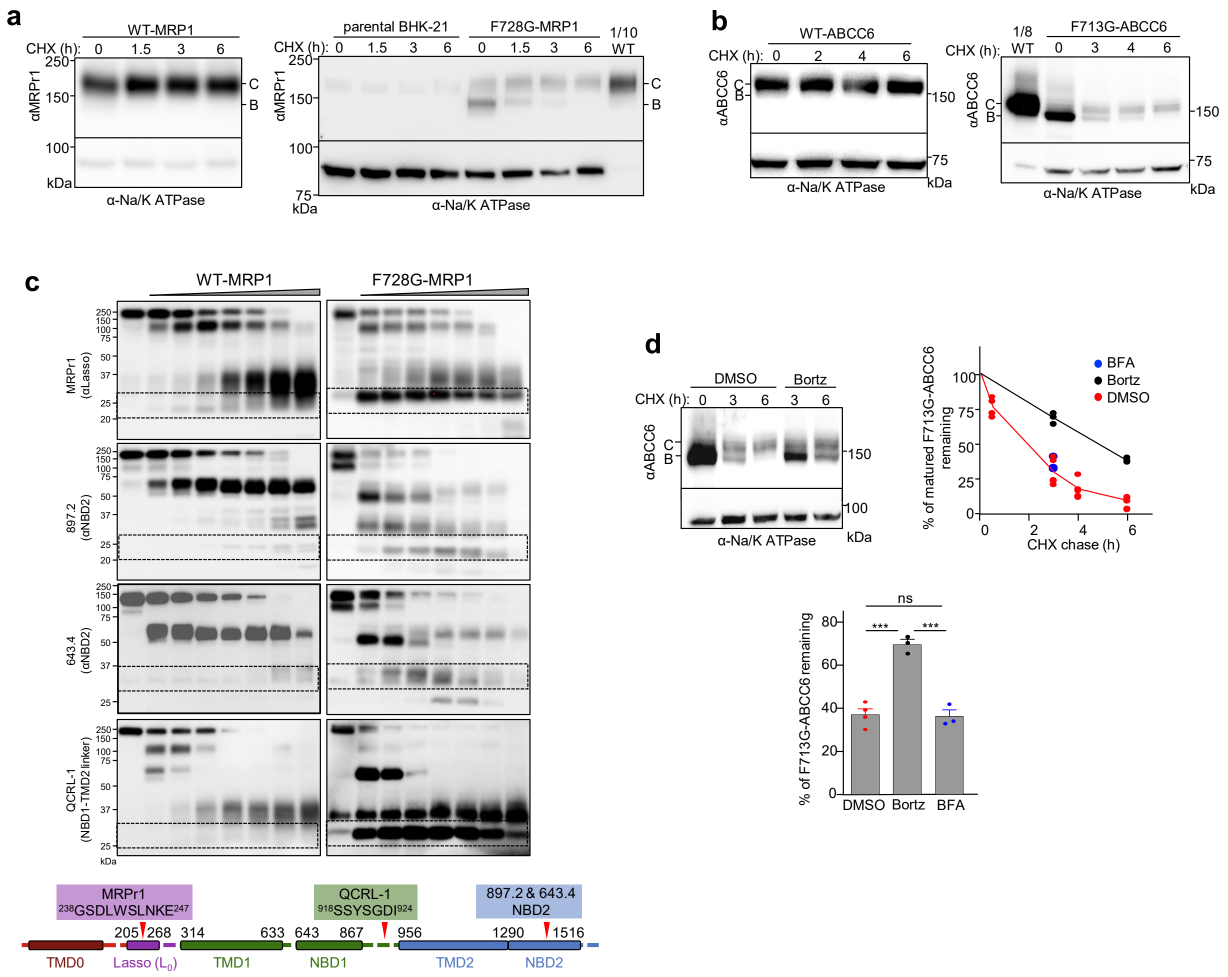


**Supplementary Figure 7. Uncoupling of the NBD1-TMD2 interface compromises ER processing and coupled domain folding of the human MRP1 and ABCC6.**

**a)** The metabolic turnover of complex-glycosylated (C-band) WT-MRP1 (left panel) and F728G-MRP1 was determined by CHX chase (150 µg/ml) and quantitative immunoblotting by loading equal amount of BHK-21 cell lysates. Only 10 % of the WT-MRP1 lysate was used for reference. The heterologous expression of MRP1 was corrected with the amount of endogenously expressed MRP1 in the quantitative analysis. **b)** The metabolic turnover of complex-glycosylated WT-ABCC6 and F713G-ABCC6 was determined by CHX-chase and quantitative immunoblotting after pre-clearing the immature forms with 3 h of CHX chase. Means ± S.E.M., n = 3, biological replicates. **c)** The impact of NBD1-CL7 interface uncoupling by the F728G mutation on the conformational stability of the WT-MRP1 NBD1, NBD2, TMD1 and TMD2 was probed by limited trypsinolysis and immunoblotting with domain-specific monoclonal antibodies (lower panel). Microsomes were isolated from BHK-21 cells expressing the indicated MRP1 variants and digested at increasing trypsin concentrations (0, 1, 3, 10, 30 100, 300 and 1000 μg/ml). Representative of two independent experiments. Based on the Ab epitope location/specificity and the proteolytic fragment size, the dashed squares indicate the following domains and their fragments accumulation and/or elimination upon of trypsinolysis: MPRr1 Ab, TMD0 and/or TMD1; 897.2 & 643.4 Abs, NBD2; QCRL-1 Ab, NBD1 and/or TMD2. **d)** The effect of proteasome (200 nM Bortezomib) and the vesicular ER-export (500 nM BrefeldinA) inhibitor on the misfolded core-glycosylated F713G-ABCC6 ER-degradation. The core-glycosylated F713G-ABCC6 (B-band) disappearance was measured by CHX-chase in the presence and absence of inhibitors (top left panel) by quantitative immunoblotting and expressed as the percentage of the initial amount as a function of the chase time after 15 min of pre-treatment with inhibitors. The effect of inhibitors was determined following the indicated CHX-chase (top right and lower panels). Means ± S.E.M, n = 3, biological replicates. Only the proteasome but not the ER-export inhibition impedes the F713G-ABCC6 degradation. Source data are provided as a Source Data file.

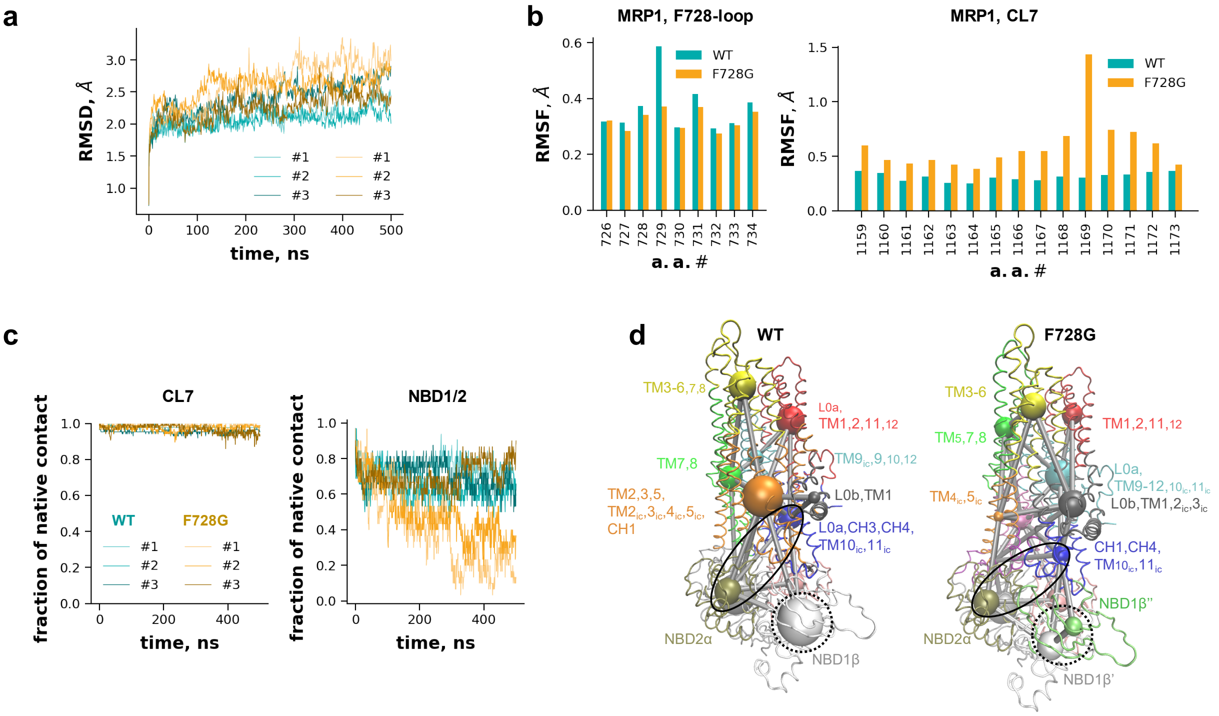


**Supplementary Figure 8. Uncoupling the NBD1-CL7(TMD2) interface likely imposes a kinetic barrier on the MRP1 cooperative domain folding landscape and stability.**

**a)** The Cα RMSD of each frame from the initial structure was calculated and indicated a sufficient equilibration process for this large protein and system (n = 3-3). **b)** RMSF values were calculated from merged trajectories (frames = 15,000) and shown for the F728-loop and CL7. **c**) Fraction of native contacts of CL7 residues and native contacts between NBD1 and NBD2 are plotted (frames = 15,000). **d**) Dynamic networks are shown in their structural context. Dynamic communities of WT- and F728G-MRP1 are represented with colored spheres and their couplings are indicated with gray sticks. The size of spheres and thickness of connecting sticks correlate with the number of amino acids within a community and level of coupling between communities, respectively. Smaller letters indicate that only a smaller fraction of the region contributes to the community. Black dotted circles highlight the allosteric decoupling of residues within the NBD1 β-subdomain. Black circles indicate the abolished allosteric communication between the L0/Lasso region and the α-subdomain of NBD2. Source data are provided as a Source Data file.

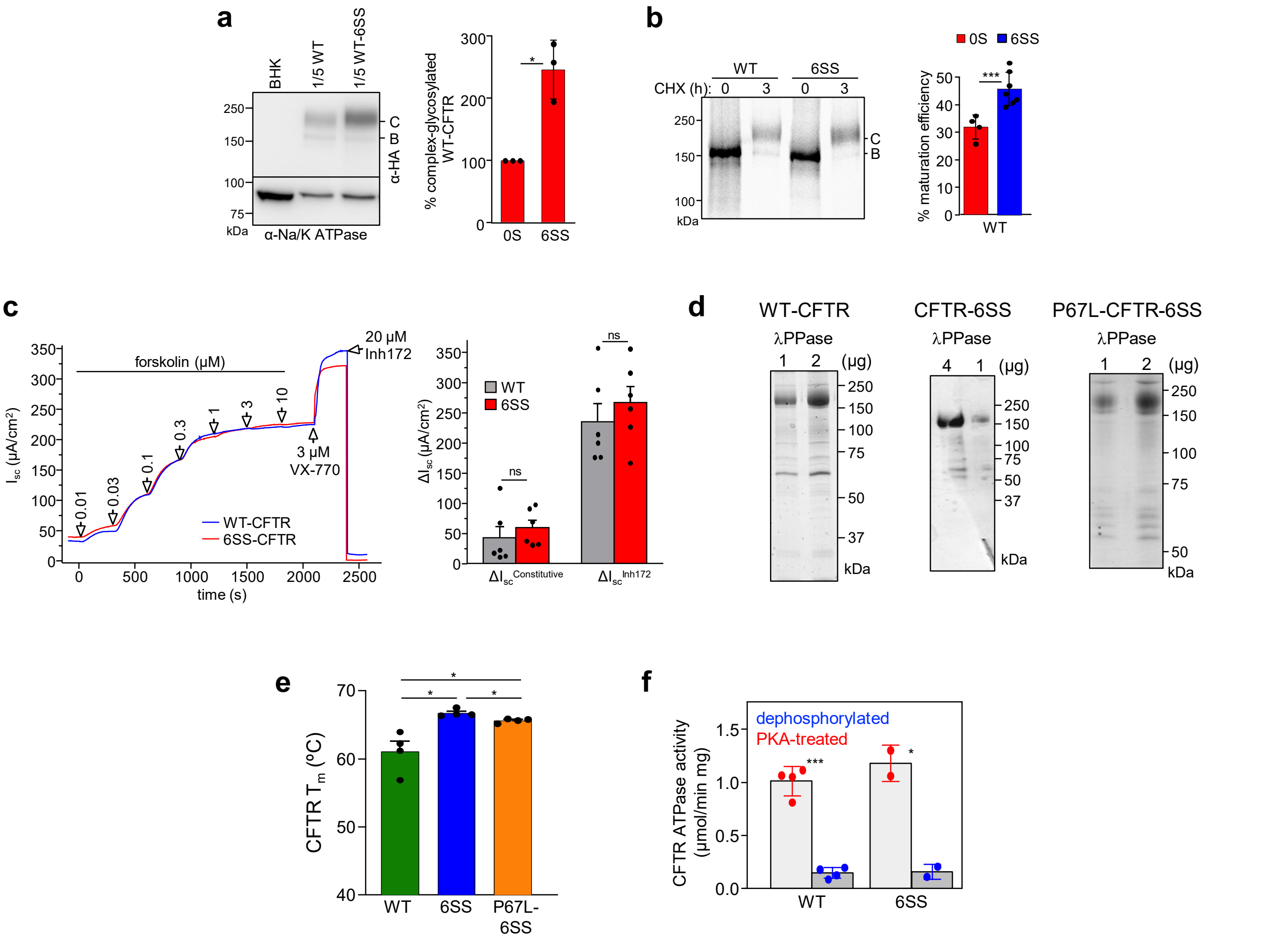


**Supplementary Figure 9. Characterization of the purified WT-, 6SS-, and P67L-6SS-CFTR channel stability and ATPase activity and their PKA-dependent activation in human bronchial epithelia (CFBE14o-) cells.**

**a)** The 6SS suppressors augment the complex-glycosylated WT-CFTR expression in BHK-21 cells, determined by quantitative immunoblotting with anti-HA Ab. Equal amounts of cell lysate were loaded. Na/K-ATPase detection was used as loading control. Means ± S.E.M., n = 3-4, biological replicates. **b)** Post-translational conformational maturation efficiency of core-glycosylated CFTR variants was measured by phosphorimage analysis after $^{35}$S-methionine and $^{35}$S-cysteine pulse-labelling (15 min) and chase (3 h) of BHK-21 cells[5]. Data are means ± S.E.M., n = 4. **c)** Functional characterization of the hyperstabilized WT- and 6SS-CFTR by short circuit current ($I_{sc}$) measurements. CFTR was activated by sequential addition of forskolin at the indicated concentration and the VX-770 gating potentiator (3 μM), followed by CFTR inhibition with $CFTR_{inh172}$ (Inh172, 20 μM) in stably transfected CFBE14o- monolayers with basolateral-to-apical chloride gradient (left panel).

Representative analogue traces. Quantification of the $CFTR_{inh172}$-inhibited currents before ($\Delta I_{sc}^{constitutive}$) and after ($\Delta I_{sc}^{Inh172}$) forskolin stimulation. Means ± S.E.M., n=3 comprising two technical replicates of each. **d)** SDS-PAGE analysis of purified CFTR variants treated by λPPase to dephosphorylate the channel, respectively. **e)** CFTR thermal stability was monitored by DSF using Bodipy-FL-Cysteine as the reporter dye. $T_m$ values were calculated by the Boltzmann equation. Means ± S.E.M., n = 4. **f)** ATP hydrolysis of CFTR was determined by the NADH-coupled assay as described in Methods. The rate of ATP hydrolysis was calculated from the equation: ATPase rate [$min^{-1}$] = $-dA_{340}$ [OD/min] × $K_{path}^{-1}$ × $moles^{-1}$, where $K_{path}$ is the molar absorption coefficient for NADH for a given optical path length. Rates were corrected for the background NADH decomposition of controls, containing no ATPase. Means ±S.D., n = 2-3 measurements from two independently purified protein preparations. Source data are provided as a Source Data file.

**a**

| CFTR domain | coverage (%) | # of peptides | mean peptide length (a.a.) |
|---|---|---|---|
| TMD1 | 81.6 | 152 | 8.4 |
| TMD2 | 56.1 | 92 | 10.2 |
| NBD1 | 100 | 208 | 9.4 |
| NBD2 | 100 | 206 | 10.8 |
| RD | 100 | 106 | 11.3 |
| **Total** | **84.2** | **764** | **9.9** |

**c**

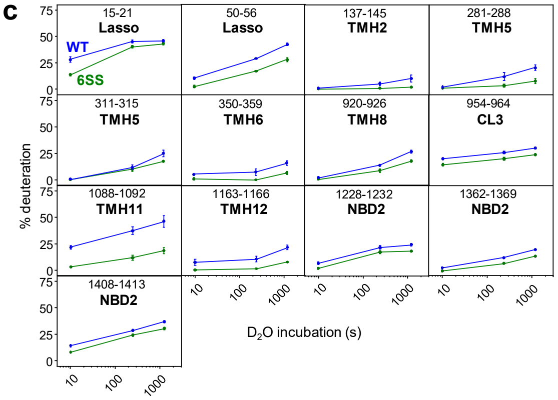


**b**

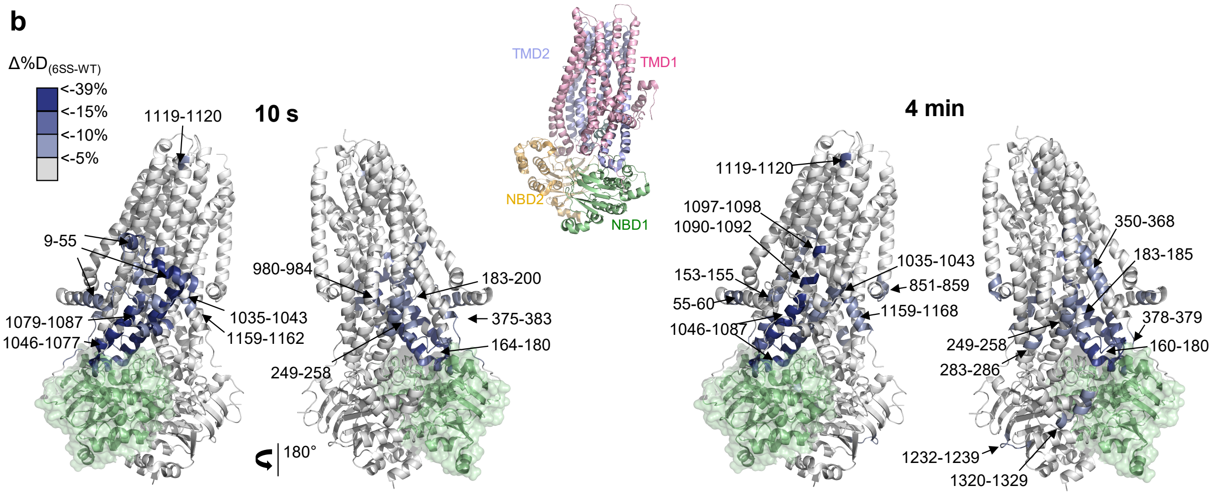


**Supplementary Figure 10. The 6SS-induced backbone dynamic stabilization of the NBD1 allosterically propagates to the TMD1, TMD2 and NBD2 shown by differential HDX kinetics of purified WT- and 6SS-CFTR.**

**a)** The peptide sequence coverage produced by on-line tandem nep2-pepsin columns digestion of WT-CFTR at 18 °C. The percentage of peptide coverage, the number of peptides and the mean peptide length for each domain and full-length CFTR are summarized. **b)** The differential deuteration (Δ%D = $\%D_{6SS}$ - $\%D_{WT}$) of TMD1/2 and NBD2 peptides was displayed on the CFTR cryo-EM structure (PDB:6MSM) after 10 s and 4 min $D_2O$ incubation at 37 °C. For clarity, only the Δ%D of TMD1/2 and NBD2 were mapped. The NBD1 is colored in green. **c)** Deuteration kinetics of selected peptides of purified WT- and 6SS-CFTR at the indicated topological location. Means ± S.E.M., n=3, technical replicates. Deuteration kinetics for the entire purified WT- and 6SS-CFTR is shown in Supplementary Fig. S11. Source data are provided as a Source Data file.

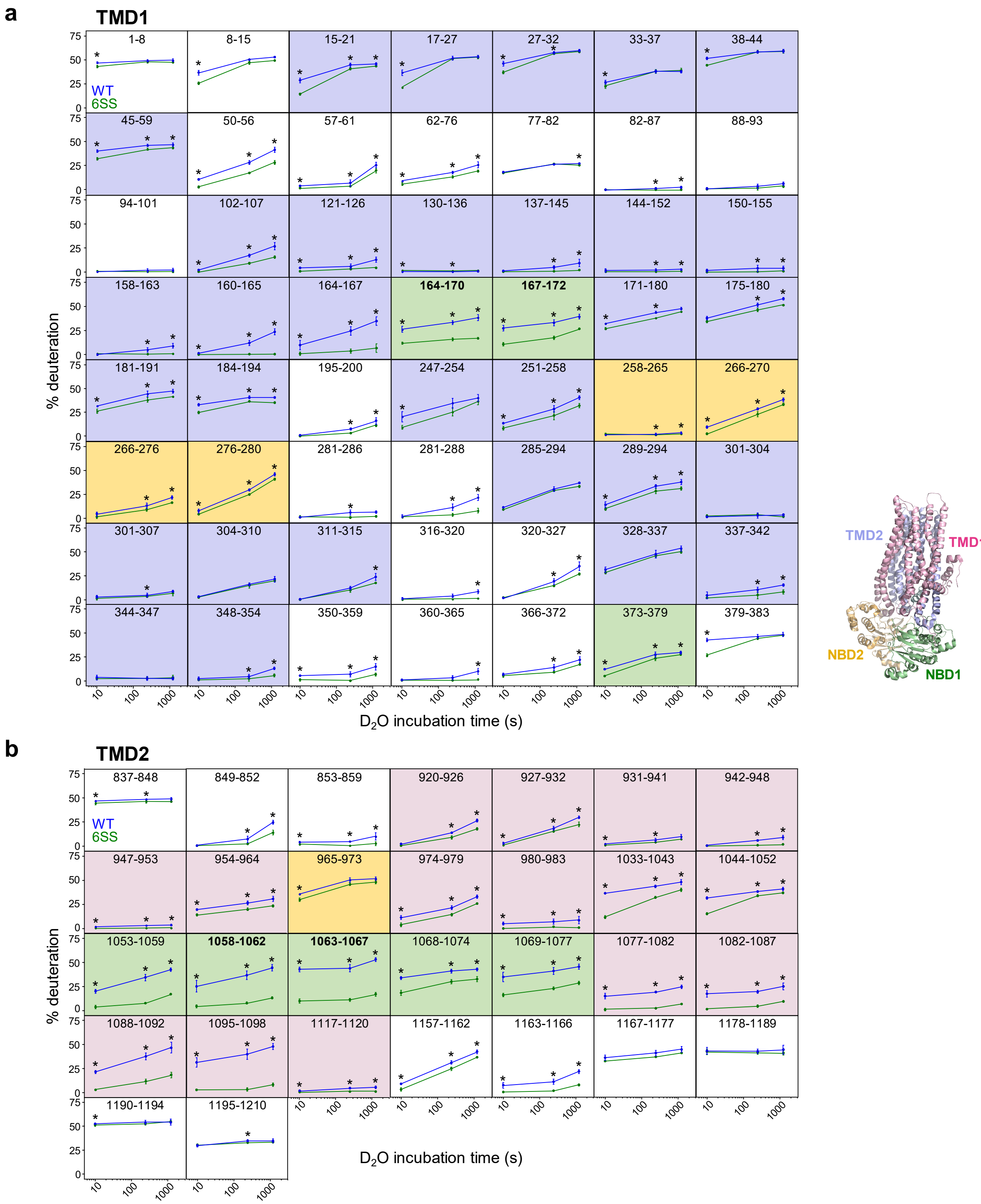
a
TMD1
1-8
8-15
15-21
17-27
27-32
33-37
38-44
45-59
50-56
57-61
62-76
77-82
82-87
88-93
94-101
102-107
121-126
130-136
137-145
144-152
150-155
158-163
160-165
164-167
164-170
167-172
171-180
175-180
181-191
184-194
195-200
247-254
251-258
258-265
266-270
266-276
276-280
281-286
281-288
285-294
289-294
301-304
301-307
304-310
311-315
316-320
320-327
328-337
337-342
344-347
348-354
350-359
360-365
366-372
373-379
379-383
WT
6SS
% deuteration
D2O incubation time (s)
TMD2
TMD1
NBD2
NBD1
b
TMD2
837-848
849-852
853-859
920-926
927-932
931-941
942-948
947-953
954-964
965-973
974-979
980-983
1033-1043
1044-1052
1053-1059
1058-1062
1063-1067
1068-1074
1069-1077
1077-1082
1082-1087
1088-1092
1095-1098
1117-1120
1157-1162
1163-1166
1167-1177
1178-1189
1190-1194
1195-1210
WT
6SS
% deuteration
D2O incubation time (s)

**c** **NBD1**

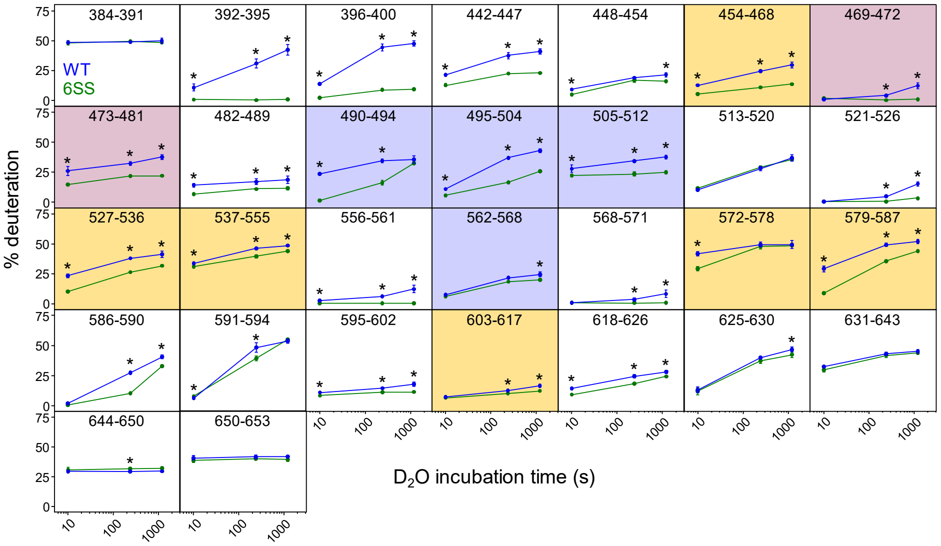


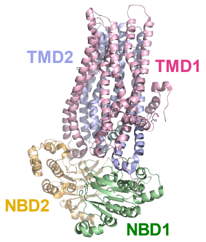


**d** **NBD2**

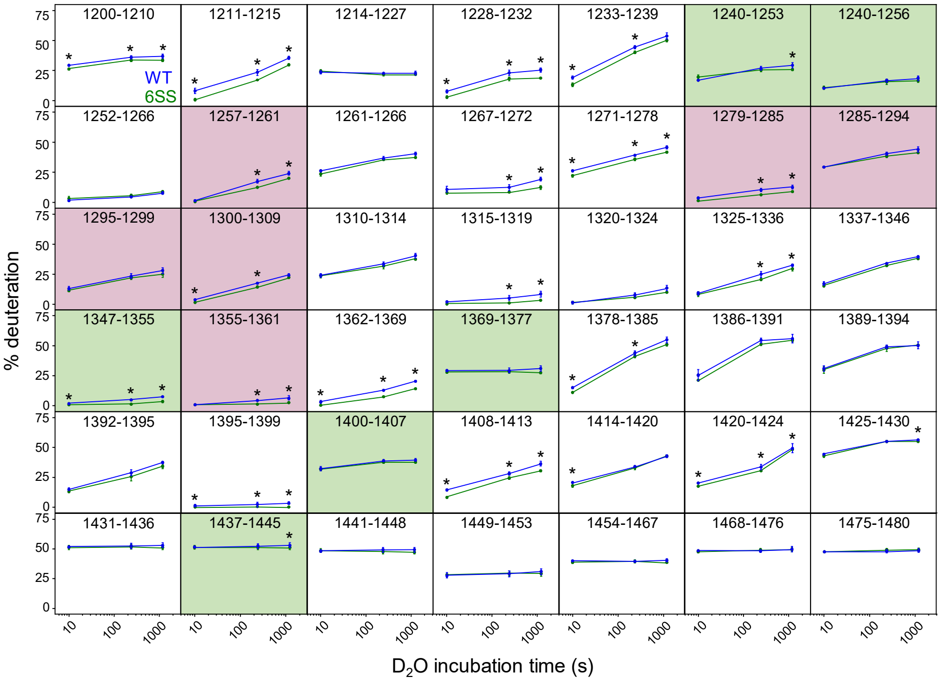


**e** **RD**

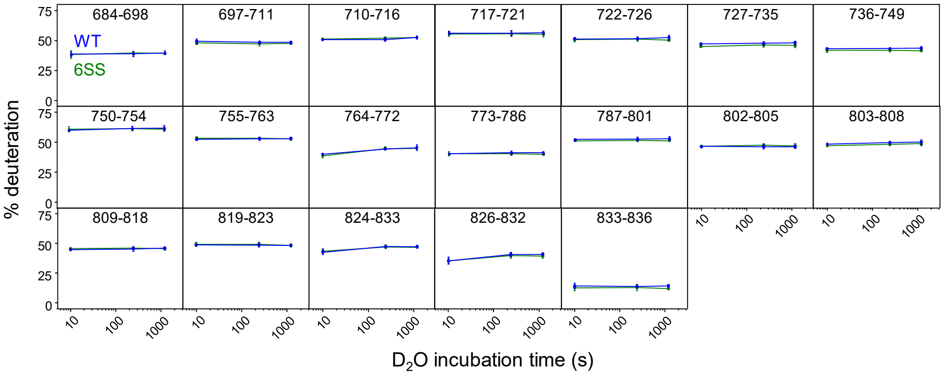

**Supplementary Figure 11. Backbone conformational dynamics of purified WT- and 6SS-CFTR by HDX-MS.**

The deuteration time course of WT- and 6SS-CFTR at 37 °C are depicted by representative peptides and presented domain wise: **a)** TMD1, **b)** TMD2, **c)** NBD1, **d)** NBD2 and **e)** regulatory domain (RD) (means ± S.D., n = 3, technical replicates). Significant difference was determined by a two-tailed Student's *t*-test ($P < 0.05$ (*). Exact P values of individual peptides are provided in the Source Data file. The background color of individual peptide indicates the predicted domain interaction (according to the inset), based on the full-length CFTR cryo-EM structure (PDB:6MSM) coloring. The entire deuteration data sets are included in the Source Data file.

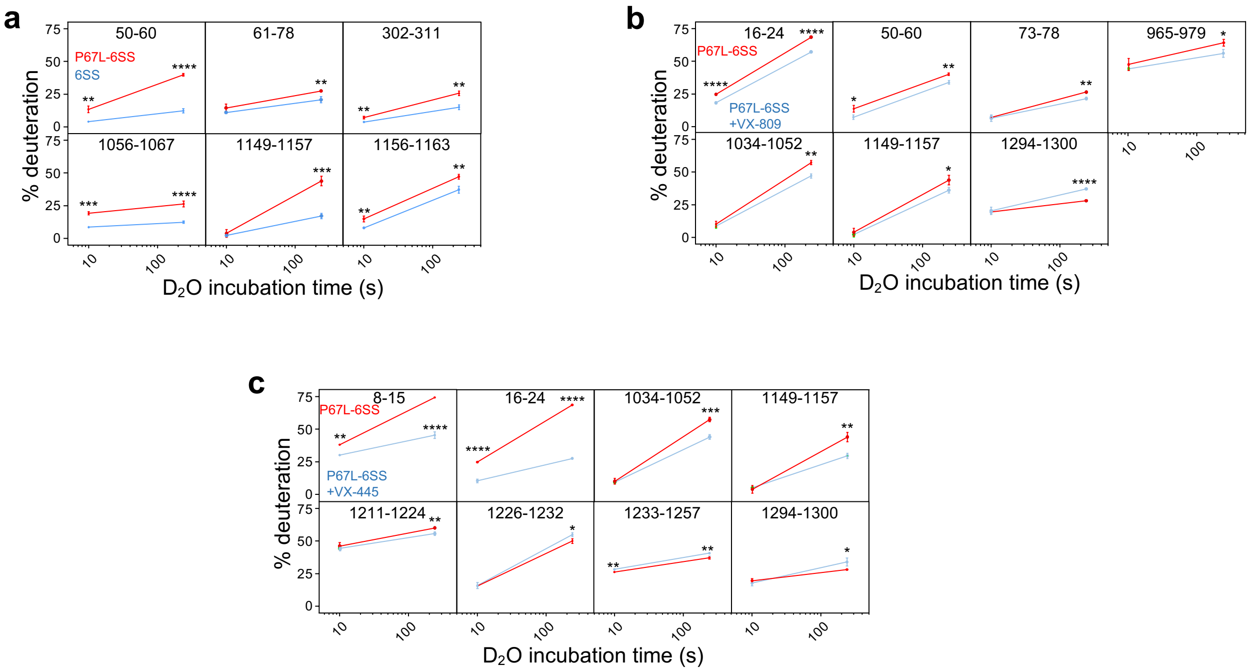


**Supplementary Figure 12. The effects of VX-809 or VX-445 on the backbone conformational dynamics of purified P67L-CFTR-6SS.**

The deuteration time course of **a)** CFTR-6SS and P67L-CFTR-6SS, **b)** P67L-CFTR-6SS in the absence and presence of VX-809 and **c)** P67L-CFTR-6SS in the absence and presence of VX-445 at 37 °C are depicted by representative peptides (means ± S.D., n = 3, technical replicates). Significant difference was determined by unpaired two-tailed *t*-test (P < 0.05 (*)). The exact P values are provided in a Source Data file. The corresponding bar plots after 4 min incubation are shown in Fig 6. Source data are provided as a Source Data file.

**Supplementary Table 1.** Conformational maturation efficiency of CFTR variants. The indicated maturation efficiency of CFTR variants was measured by radioactive pulse-chase experiments as described in Methods. The maturation efficiency was corrected for CFTR variants that display accelerated turnover of their complex-glycosylated forms. The turnover of the complex-glycosylated CFTR was determined by cycloheximide (CHX) chase and immunoblotting (Supplementary Fig. 1e) as described in Methods.

| **CFTR construct** | maturation efficiency (%) (means ± S.E.) | corrected maturation efficiency (%) | N | CFTR variant maturation efficeincy % (mean ± S.E.) with corrector drugs exposed during: | | | | | | | | | | | |
|---|---|---|---|---|---|---|---|---|---|---|---|---|---|---|---|
| | | | | depletion | N | pulse | N | chase | N | depletion + pulse | N | pulse + chase | N | depl + pulse + chase | N |
| WT | 32.46 ± 1.12 | | 5 | | | | | | | | | | | | |
| WT-6SS | 44.22 ± 1.81 | | 5 | | | | | | | | | | | | |
| P67L | 1.96 ± 0.38 | | 4 | | | | | 34.6 ± 5.2 | 6 | | | | | | |
| L206W | 0.67 ± 0.16 | | 4 | 3.0 ± 1.34 | 3 | 3.1 ± 1.07 | 5 | 31.7 ± 2.2 | 12 | 9.3 ± 1.8 | 3 | 34.8 ± 5.8 | 4 | 42 ± 3.79 | 3 |
| ΔF508 | 1.2 ± 0.24 | | 3 | | | | | 32.1 ± 3.6 | 6 | 6 ± 3 | 4 | 8.9 ± 1.7 | 4 | 36.2 ± 2 | 6 |
| ΔF508-6SS | 42.27 ± 1.23 | | 5 | | | | | | | | | | | | |
| F508G | 2.06 ± 0.31 | 2.3 ± 0.7 | 5 | | | | | | | | | | | | |
| F508G-2PT | 20.78 ± 1.34 | | 5 | | | | | | | | | | | | |
| R170G+F508G | 1.32 ± 0.26 | 2.64 ± 0.52 | 5 | | | | | | | | | | | | |
| R170G+F508G-2PT | 4.02 ± 0.27 | 6.86 ± 0.55 | 5 | | | | | | | | | | | | |

**Supplementary Table 2** Expression constructs for the WT and mutant variants of isolated NBD1s and full-length CFTR, MRP1 and ABCC6.

**CFTR isolated NBD1**

**Suppressors**

| Name | Mutations | Source |
|---|---|---|
| WT | None | Rabeh et al. 2012 |
| 2PT | deletion of amino acid 404-435, S492P, A534P, I539T | *This paper* |
| 6SS | deletion of amino acid 404-435, M470V, S492P, S495P, A534P, I539T, R555K | *This paper* |

**Domain interface mutations**

| Name | Mutations | Source |
|---|---|---|
| F508G | F508G | *This paper* |
| F508G-2PT | F508G, deletion of amino acid 404-435, S492P, A534P, I539T | *This paper* |

**Full-length CFTR**

**Suppressors**

| Name | Mutations | Source |
|---|---|---|
| WT | None | Du and Lukacs, 2009 |
| 6SS | deletion of amino acid 404-435, M470V, S492P, S495P, A534P, I539T, R555K | *This paper* |

**Domain interface mutations**

| Name | Mutations | Source |
|---|---|---|
| F508G | F508G | *This paper* |
| F508G-2PT | F508G, deletion of amino acid 404-435, S492P, A534P, I539T | *This paper* |
| R170G | R170G | *This paper* |
| R170G-3S | R170G, F429S, F494N, Q637R | *This paper* |
| R170G-2PT | R170G, deletion of amino acid 404-435, S492P, A534P, I539T | *This paper* |

**CF mutations**

| Name | Mutations | Source |
|---|---|---|
| ΔF508 | deletion of F508 | Du and Lukacs, 2009 |
| P67L | P67L | *This paper* |
| P67L-6SS | P67L, deletion of amino acid 404-435, M470V, S492P, S495P, A534P, I539T, R555K | *This paper* |
| L206W | L206W | *This paper* |

**MRP1 isolated NBD1**

| Name | Mutations | Source |
|---|---|---|
| WT | None | *This paper* |
| F728G | F728G | *This paper* |

**Full-length MRP1**

| Name | Mutations | Source |
|---|---|---|
| WT | None | *This paper* |
| F728G | F728G | *This paper* |

**ABCC6 isolated NBD1**

| Name | Mutations | Source |
|---|---|---|
| WT | None | *This paper* |
| F713G | F713G | *This paper* |

**Full-length ABCC6**

| Name | Mutations | Source |
|---|---|---|
| WT | None | *This paper* |
| F713G | F713G | *This paper* |

**Supplementary Table 3.** Distribution of the distance between NBDs calculated from the MD simulation trajectories from the 450-500 ns interval.

**WT-CFTR**

| | Site-1* | Site-2 |
|---|---|---|
| #1*** | 13.0 (1.0) | 11.3 (1.3) |
| #2 | 10.5 (0.2) | 9.9 (0.2) |
| #3 | 12.3 (0.2) | 12.4 (0.5) |
| #4 | 12.1 (0.2) | 9.9 (0.2) |
| #5 | 12.3 (0.3) | 9.9 (0.2) |
| #6 | 12.0 (0.4) | 12.7 (0.4) |
| #1-#6 | 12.0 (0.9) | 11.0 (1.3) |

Distribution**

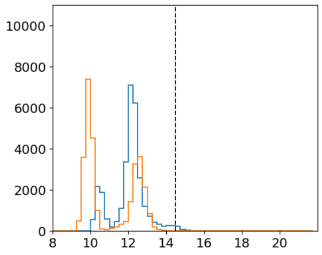

**R170G-CFTR**

| | Site-1 | Site-2 |
|---|---|---|
| #1 | 12.2 (0.2) | 13.1 (0.3) |
| #2 | 12.2 (0.2) | 10.0 (0.2) |
| #3 | 12.2 (0.2) | 10.2 (0.2) |
| #4 | 10.6 (0.2) | 9.8 (0.3) |
| #5 | 10.6 (0.2) | 10.1 (0.2) |
| #6 | 10.7 (0.2) | 12.5 (0.3) |
| #1-#6 | 11.4 (0.8) | 11.0 (1.3) |

Distribution

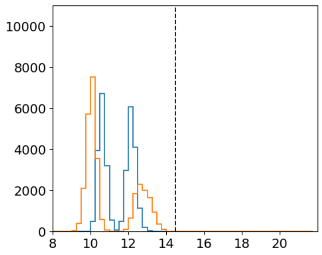

**F508G-CFTR**

| | Site-1 | Site-2 |
|---|---|---|
| #1 | 10.5 (0.2) | 9.9 (0.2) |
| #2 | 15.1 (0.4) | 9.9 (0.2) |
| #3 | 12.1 (0.2) | 9.7 (0.3) |
| #4 | 19.2 (1.1) | 12.6 (0.6) |
| #5 | 10.6 (0.2) | 9.8 (0.2) |
| #6 | 11.9 (0.2) | 14.0 (0.6) |
| #1-#6 | 13.2 (3.1) | 11.0 (1.7) |

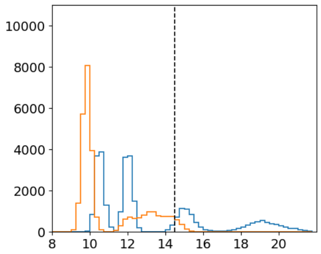

**P67L-CFTR**

| | Site-1 | Site-2 |
|---|---|---|
| #1 | 16.3 (1.0) | 12.3 (0.3) |
| #2 | 10.5 (0.2) | 9.9 (0.2) |
| #3 | 12.1 (0.2) | 10.0 (0.3) |
| #4 | 12.1 (0.2) | 10.0 (0.2) |
| #5 | 10.6 (0.2) | 12.5 (0.3) |
| #6 | 12.5 (0.3) | 10.0 (0.2) |
| #1-#6 | 12.3 (2.0) | 10.8 (1.2) |

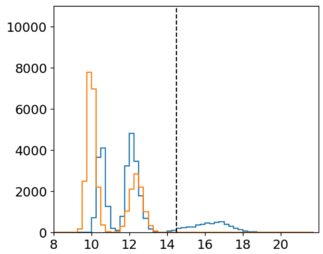

**F508G-6SS-CFTR**

| | Site-1 | Site-2 |
|---|---|---|
| #1 | 13.2 (0.4) | 9.8 (0.3) |
| #2 | 12.1 (0.4) | 10.1 (0.2) |
| #3 | 13.0 (0.3) | 9.7 (0.2) |
| #4 | 10.6 (0.4) | 10.0 (0.5) |
| #5 | 10.6 (0.2) | 9.9 (0.3) |
| #6 | 12.3 (0.3) | 10.2 (0.2) |
| #1-#6 | 12.0 (1.1) | 10.0 (0.3) |

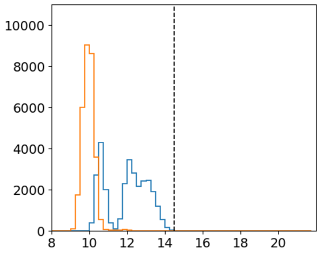

**P67L-6SS-CFTR**

| | Site-1 | Site-2 |
|---|---|---|
| #1 | 12.2 (0.2) | 9.9 (0.2) |
| #2 | 10.4 (0.2) | 10.4 (0.2) |
| #3 | 12.2 (0.3) | 9.9 (0.2) |
| #4 | 10.6 (0.2) | 10.1 (0.2) |
| #5 | 15.6 (0.7) | 10.1 (0.2) |
| #6 | 12.4 (0.3) | 10.0 (0.2) |
| #1-#6 | 12.2 (1.7) | 10.1 (0.3) |

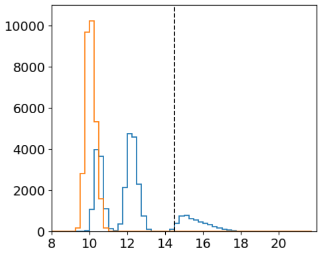

**WT-MRP1**

| | Site-1 | Site-2 |
|---|---|---|
| #1 | 10.6 (0.2) | 10.0 (0.2) |
| #2 | 10.7 (0.2) | 10.0 (0.2) |
| #3 | 13.4 (0.5) | 10.1 (0.2) |
| #1-#3 | 11.6 (1.3) | 10.0 (0.2) |

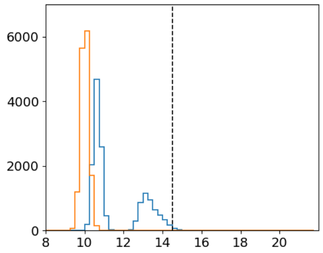

**F728G-MRP1**

| | Site-1 | Site-2 |
|---|---|---|
| #1 | 17.3 (1.1) | 13.1 (0.5) |
| #2 | 12.4 (0.7) | 10.1 (0.2) |
| #3 | 10.4 (0.2) | 10.1 (0.2) |
| #1-#3 | 13.4 (3.0) | 11.1 (1.5) |

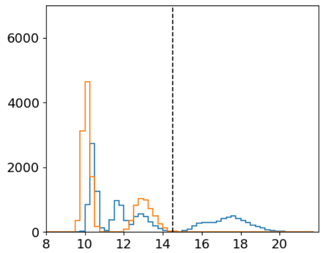

* mean (standard deviation) of distance between Walker A and opposite signature motif (for CFTR Site-1: 464/1348 and Site-2: 1250/550, for MRP1 Site-1: 684/1431 and Site-2: 770/1332)

** frequency (y axis) versus distance (Å) distribution of CFTR and MRP1 from six (#1-#6) and three (#1-#3) trajectories, respectively; vertical dashed line is 14.5 Å cutoff for closed/open ATP-binding sites[6]

*** trajectory #i
